# Changes in functional connectivity preserve scale-free neuronal and behavioral dynamics

**DOI:** 10.1101/2023.09.13.557619

**Authors:** Anja Rabus, Davor Curic, Victorita E. Ivan, Ingrid M. Esteves, Aaron J. Gruber, Jörn Davidsen

## Abstract

Does the brain optimize itself for storage and transmission of information and if so, how? The critical brain hypothesis is based in statistical physics and posits that the brain self-tunes its dynamics to a critical point or regime to maximize the repertoire of neuronal responses. Yet, the robustness of this regime, especially with respect to changes in the functional connectivity, remains an unsolved fundamental challenge. Here, we show that both scale-free neuronal dynamics and self-similar features of behavioral dynamics persist following significant changes in functional connectivity. Specifically, we find that the psychedelic compound ibogaine that is associated with an altered state of consciousness fundamentally alters the functional connectivity in the retrosplenial cortex of mice. Yet, the scale-free statistics of movement and of neuronal avalanches among behaviorally-related neurons remain largely unaltered. This indicates that the propagation of information within biological neural networks is robust to changes in functional organization of sub-populations of neurons, opening up a new perspective on how the adaptive nature of functional networks may lead to optimality of information transmission in the brain.

---

The brain integrates multi-sensory information in order to generate appropriate responses to a wide range of environmental stimuli. This information must be flexibly routed within and between brain regions as part of this process. Although the mechanisms of information routing remain unclear, a common working theory is that information is routed via transiently activated patterns of functionally connected neuronal populations [1]. A flexible functional network facilitates formation of new activation patterns and adaptation of existing ones, which enables the brain to generate sequences of appropriate patterned activity even to novel stimuli. Despite significant headway made in studying the coordinated activity of neuronal populations, the role of functional connectivity (FC) in the mechanism of information routing and neural dynamics remains poorly understood.

The critical brain hypothesis is a framework based in statistical physics to account for the properties of collective neuronal activity and information processing in the brain [2–4]. It posits that quiet wakefulness is characterized by a self-organized critical state in which the repertoire of possible neuronal activity patterns is maximized [5–8]. Hallmarks of such a non-equilibrium phase transition include scale-free cascades of causally connected neuronal activity (dubbed neuronal avalanches) [9–11], and the presence of long-range spatial [12] and temporal [13–15] correlations, all of which have been observed *in vivo* [9, 16], *in vitro* [17, 18], and across species [19–23] and spatial scales [14, 23–27]. This suggests the existence of some universal mechanism or process that imbues brain networks with the flexibility required for optimization of information storage and transmission. The repertoire of accessible neuronal activation states is maximized at criticality, allowing network dynamics to adapt both to situations that require immediate attention (e.g. task or rapidly-changing situation) and to long-term stimuli (e.g. stress).

Although critical dynamics are thought to be important for information processing, the explicit relationship of this ensemble property to information routing, FC and behavior is not currently known. E.g., to what extent are topological changes in FC reflected in signatures of criticality? Certainly, large changes in neural excitability result in observable changes to critical dynamics, such as the out-right loss of scale-free statistics under surgical-plane anesthesia [7, 10, 28, 29], or changes in the universality class brought about by pathological neural development in cultures [30], among others [31–33]. While loss of criticality due to changes in excitability can be tied to changes in the overall degree of correlations among neurons [7], the relationship to specific topological changes in FC, information routing and behavior has remained elusive. It is an open question how perturbations to FC affect signatures of criticality under normal physiological conditions.

To tackle this challenge, we studied the robustness of neuronal and behavioral dynamics to changes in FC in mouse retrosplenial cortex (RSC), an area of the brain well positioned to integrate sensory, mnemonic, and cognitive information by virtue of its strong connectivity with the hippocampus, medial prefrontal cortex, and primary sensory cortices. As shown in [34–37], a subset of RSC neurons increase neural activity during movement, while another decreases activity. In the present study, we perturbed these functional networks using the psychedelic drug ibogaine. Ibogaine has been shown to increase mean excitability [38] but its effects on network dynamics at the single-cell level are not well understood. We compare the resulting changes in behavior and neuronal dynamics to a control case in which we administered saline. We find that while FC is significantly altered under ibogaine, the signatures of criticality in both behavior and neuronal dynamics are not. Observing the same behavioral output and the same overall neuronal dynamics at the network level under changes to FC is consistent with the claim that information can be routed via more than one sequence of transiently co-activating neuronal populations [39]. These results offer a new perspective on the relationship between the adaptive nature of FC and optimality of information transmission in the brain and also support the recently proposed connection between scale-free behavioral and neuronal dynamics [36].

## Experimental setup

Head-fixed mice were free to locomote or remain still on a passive treadmill belt [34]. Neural activity of several hundred neurons was recorded with 2-photon imaging of a genetically-encoded activity-dependent fluorescent sensor. Activity and belt movement was recorded for 10 minutes before administering either ibogaine or saline via intraperitoneal injection. Following a 10-minute period for the drug to take effect, neural activity and treadmill belt movement was again recorded for 10 minutes. An additional 10-minute recording was taken 30 minutes after injection. For more details on experiments and data collection, see Sec. S1A.

## Functional connectivity

Mice spontaneously move the treadmill in episodes separated by brief rest periods. We classified each neuron according to its change in neural activity as animals transitioned from stationary to moving on the treadmill (see Sec. S1B for more details). As the animal transitioned from stationary to moving, some neurons increased activity, whereas others decreased it. We refer to these as up-regulated and down-regulated, respectively. In contrast, neuron activity was largely homogeneous across all neurons when the animal was not moving, i.e., resting. Fig. 1(a) shows an example of the activity of up-regulated and down-regulated neurons with track speed. Neurons belonging to either up- or down-regulated populations are referred to as *classified*, all others are referred to as *unclassified* (see also Table S1).

**FIG. 1.**
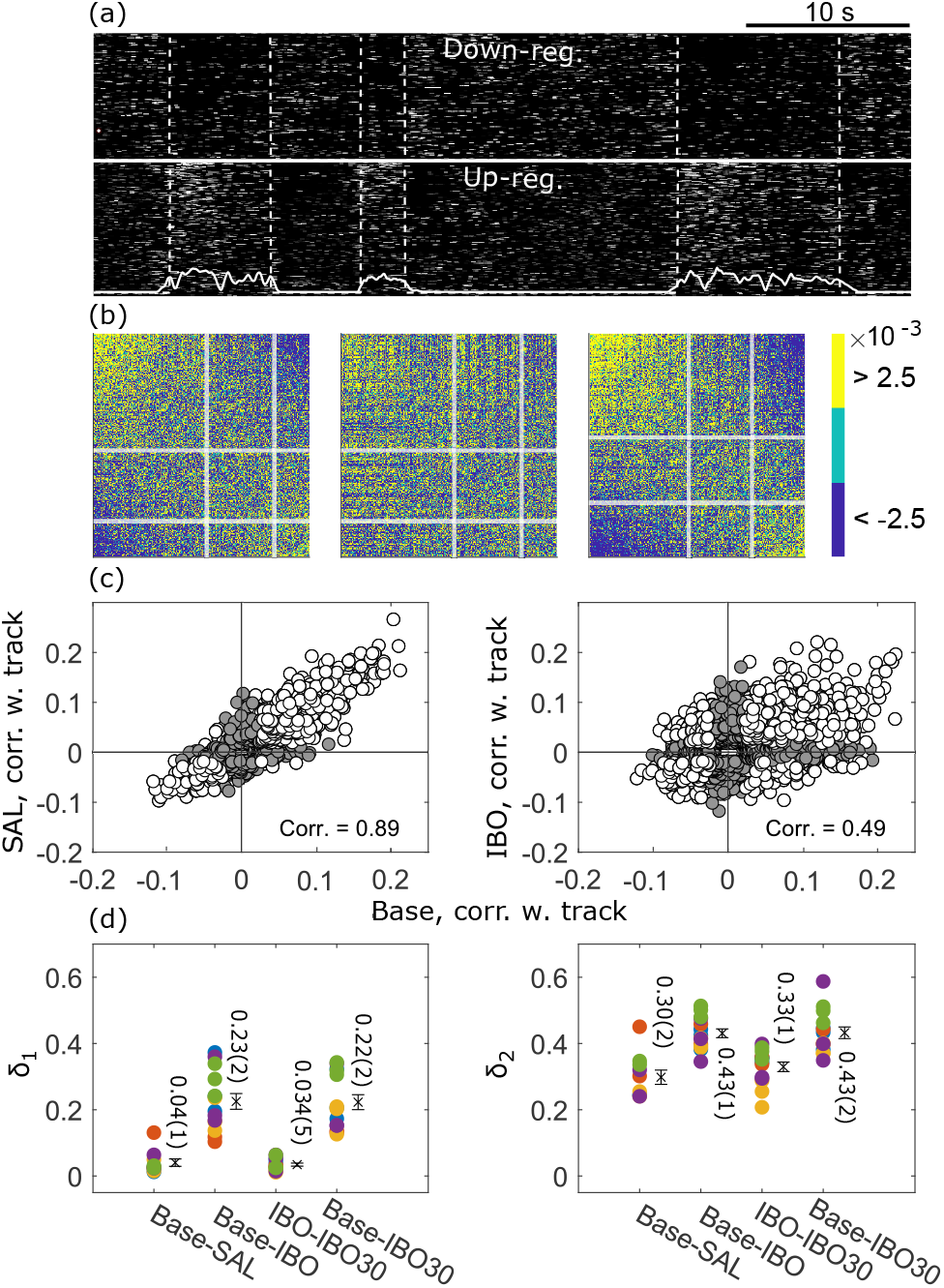
Changes in FC. (a) Activity of up- and down-regulated neurons, during moving and resting periods (separated by white dashed curve). The track speed is overlaid (solid white curve). (b) Examples of pairwise neuron correlations matricies: (Left) Ordered from most correlated to most anti-correlated with track speed for a baseline recording. White lines separate classified from unclassfied neurons. (Middle) For the subsequent ibogaine recording, preserving the order of the neurons in the left panel. (Right) For the same ibogaine recording but re-ordered to account for changes in the correlations with track speed. (c) Track speed correlation of all neurons across all post-administration recordings (saline (SAL) left, ibogaine (IBO) right) plotted against their respective baseline (Base). Cells classified in *both* baseline and post-administration recordings are represented by hollow circles. Grey circles represent cells which were unclassified in at least one recording. (d) Percentage of neurons that changed classification category from baseline to saline, baseline to ibogaine, ibogaine to 30 minutes post-ibogaine (IBO30) and baseline to IBO30. Colors represent different mice, and crosses with errorbars represent the mean and standard error. Left: Percentage of neurons that switched classification category (*δ*_1_). Right: Percentage of all neurons that were classified at least once and switched (*δ*_2_).

FC can be quantified by pairwise correlations among neurons. Fig. 1(b) illustrates this in a baseline recording (i.e., prior to drug injection) and the subsequent ibogaine recording. The pairwise correlation matrix is strongly altered after ibogaine injection but not after saline injection (Fig. S1). This phenomenon is observed in all recordings; the mean absolute matrix distance between baseline and ibogaine (0.0134(5)) is statistically greater than between baseline and saline (0.0110(4)) based on a Kolmogorov-Smirnov (KS) test, with p=0.01, persistent 30-minutes after ibogaine administration (Fig. S2). Yet, ibogaine preserves certain topological features, such as the strength of neuron-neuron correlations and weighted degree distribution of the pairwise correlation matrix (Fig. S3). This is supported by the visual similarity of the left and right panels of Fig. 1(b).

The changes in FC can be further quantified. Figure 1(c) shows that correlations between neuron activity and track speed are significantly altered under ibogaine in comparison to saline, suggesting that ibogaine modulates neuronal activation patterns in the RSC beyond what can be explained by differences in motor output. A quantification of activity pattern modulation is shown in Fig. 1(d). Circles represent the percentage of neurons that changed classification among up- and down-regulated after injection. Of the neurons that were classified in both recordings (left), 23 ± 2% switched between populations after administration of ibogaine, compared to 4 ± 1% under saline. This behavior persists 30-minutes post-ibogaine administration. Of the neurons that were classified in at least one of the two recordings (right), 43 ± 1% were re-assigned in the ibogaine recording, compared to 30 ± 2% under saline, with the change persisting at least 30 minutes post-ibogaine. These data reveal that FC is robustly altered following ibogaine administration, and persists for at least 30 minutes.

## Behavioral dynamics

To test the critical brain hypothesis under the influence of ibogaine and also to establish the effect of ibogaine on the moving and resting behavior, we studied the track movement data. The mice move at varying speed on the treadmill, and occasionally stop. We focused on periods of forward movement, preceded and succeeded by periods of immobility (see Sec. S1D for more details). These periods, which we term behavioral avalanches in this paper, can be characterized by their size *l*, i.e., the total distance covered, and their duration *t*, i.e., the total time elapsed during the forward movement. The behavioral avalanche size distribution, *P* (*l*), is shown in Fig. 2(a), and appears to follow a power-law, *P* (*l*) ∝ *l*^*τ*^, over at least 2 orders of magnitude, with *τ* = 0.81(5) The self-similar features of periods of forward movement are robust to changes in the threshold used to define movement events covering a range from non-behavioral motor activity to actual behavior (see Fig. S4). Mice typically stop moving before the end of one lap, possibly in preparation to receive their reward after each lap (see Sec. S1A), which explains the cut-off in behavioral dynamics at about 50cm. While this experimental limitation does not allow us to clearly test whether there is a difference between motor dynamics on shorter ranges and ballistic movement at longer ranges once a decision has been made by the animal, recent findings for experiments without reward administration suggest that there is no difference [36]. In any case, *P* (*l*) is consistent with the critical brain hypothesis if finite size effects are taken into account. This is further supported by the power-law relationship between *l* and *t*, ⟨*t*⟩(*l*) ∝ *l*^*γ*^, shown in Fig. 2(b) for all baseline recordings.

**FIG. 2.**
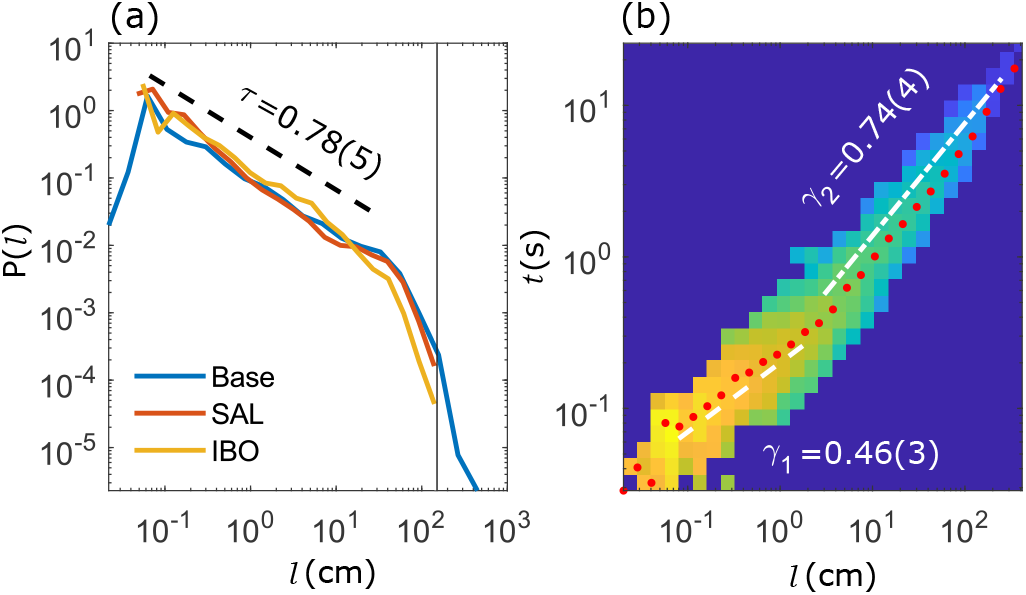
Scale-free behavioral dynamics. (a) Distribution of behavioral avalanche sizes. The exponent *τ* was estimated for the baseline recording. The vertical line at 150 cm indicates the belt length. (b) Relationship between avalanche duration and size, for all baseline recordings only. Pixel color represents the density of points, and red dots indicate the average ⟨*t*⟩ for a given *l*. The exponents *γ*_1_ and *γ*_2_ are given for two fit ranges (white dashed lines).

We observe two ranges with scaling exponents *γ*_1_ and *γ*_2_, and the transition between the regimes occurs around 1cm of track movement. For avalanches larger than 1cm, the average track speed decreased, possibly indicating the transition from sporadic to steady/regular movement. The same transition can be observed for saline and ibogaine (Fig. S5(b)), with similar values for *γ*_1_ and *γ*_2_, indicating that the relationship between avalanche size and duration is maintained under ibogaine. Due to the limited range of the behavioral avalanche duration distribution, *P* (*t*), (Fig. S5(c)) — and potentially the existence of two power-law ranges — the uncertainties are too large to get a reliable estimate of any power-law exponent *α* and to determine whether the expected scaling relation between *τ, α* and *γ* holds [40].

As suggested by Fig. 2(a), the statistical properties of behavioral avalanches were largely unaffected by saline or ibogaine (Figs. S4, S5). The only significant effect was that ibogaine reduces the largest size and duration of the behavioral avalanches — the animals have a lower mean speed (Fig. S6) and fewer long movement intervals (Fig. S5(c)). When normalizing *l* and *t* by their respective averages, ⟨*l*⟩ and ⟨*t*⟩, differences between the curves vanish (Fig. S7). This indicates that ibogaine only reduced the average size and duration of behavioral avalanches slightly, but the shape of the distributions remained form-invariant, indicating self-similarity consistent with the critical brain hypothesis. This invariance implies in particular that the ibogaine-induced changes in FC discussed above are not primarily a consequence of altered motoric output.

## Neuronal dynamics

To further test the critical brain hypothesis under the influence of ibogaine, we analyzed the statistical properties of neuronal activation. We analyzed distributions of neuronal avalanches — bursts of neuronal activity between periods of relative quiescence — for the classified populations (see Sec. S1E for more details). Specifically, we focused on avalanches conditioned on the voluntary movement behavior of the animals, i.e., resting vs moving phases as in [34], and investigated how these distributions are altered if ibogaine-induced changes to FC are or are not taken into account.

In all cases, we find size distributions resembling power laws (Figs. 3(a), S8), consistent with the critical brain hypothesis. Following Refs. [41, 42], power law exponents were estimated using maximum likelihood within the largest range [*S*_*min*_, *S*_*max*_] that supports *p >* 0.1 in a two-sample KS-test against a theoretical power-law with the same exponent (more details in Sec. S1G). The quality of the fit is measured by the dynamic range ∆ = log_10_(*S*_*max*_*/S*_*min*_). In all cases we found [*S*_*min*_, *S*_*max*_] that supported *p >* 0.1 (Fig. 3(b)), but ∆ was largest for the up-regulated group during movement, and smallest for the down-regulated group, also during movement, similar to Refs. [34, 36].

**FIG. 3.**
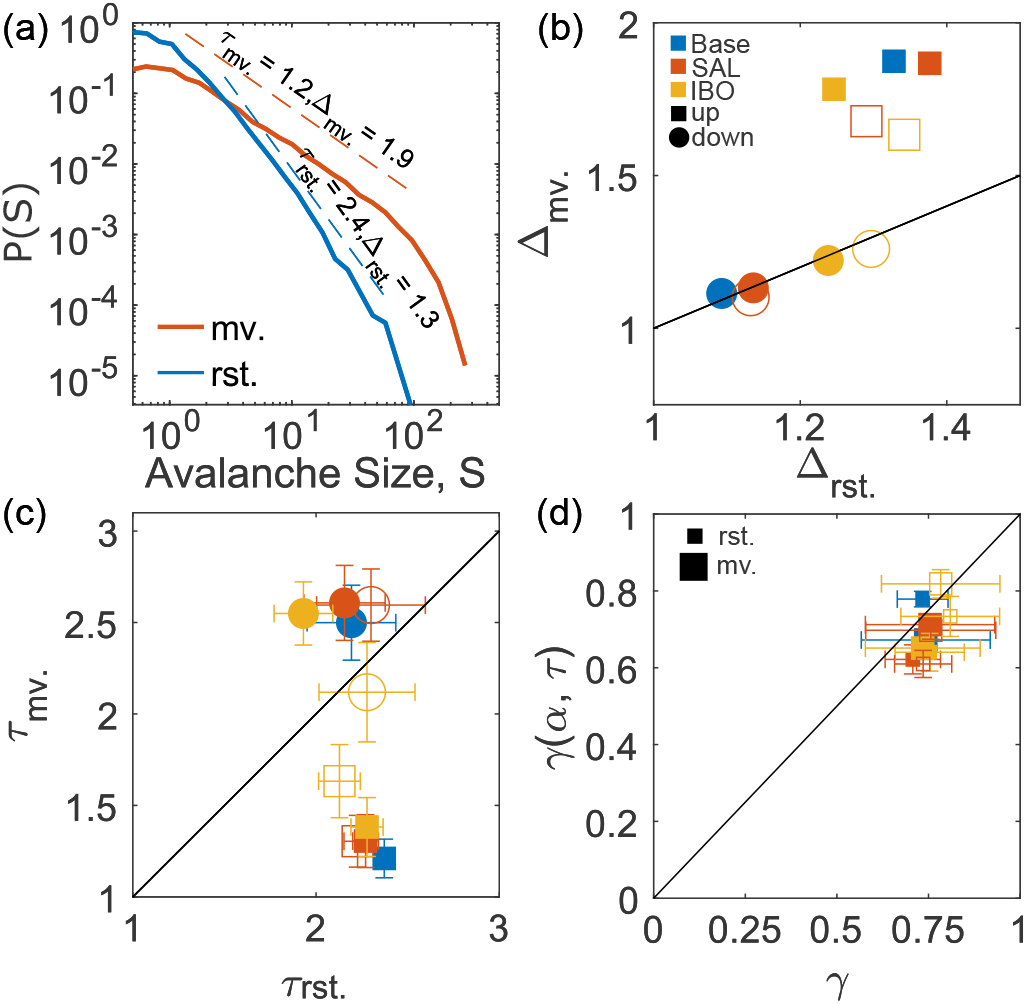
Neuronal dynamics. (a) Example of neuronal avalanche size distribution for the up-regulated sub-groups in resting (rst.) and moving (mv.) phases for baseline. The exponent *τ* and the dynamic range ∆ are also indicated. (b) The dynamic range during moving phase plotted against the resting phase equivalent, for each of the different conditions for up-(square) and down-regulated (circle) cells. In all cases, solid markers represent the result obtained when changes to FC are accounted for by re-classifying up/down neurons for each recording independently, hollow markers indicate values when these changes are not accounted for by using the previous baseline classification. *x* = *y* is shown for reference. (c) As in (b) but for the avalanche size exponents. (d) 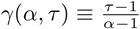 plotted vs. the directly estimated value of *γ* for different conditions for up-regulated cells. Large markers denote moving phases, small markers denote resting phases.

The estimated power-law exponents are summarized in Fig. 3(c) (see also Tables S3, S4). We find that if FC is taken into account, the estimated exponents are consistent across baseline, saline, and ibogaine recordings. During the resting phase, the exponents are statistically indistinguishable across sub-populations. In contrast, during movement, the activity of the down-regulated population is suppressed and large avalanches become rare, leading to larger exponents and a smaller ∆. Conversely, large avalanches become common in the up-regulated sub-population during movement and the estimated exponent becomes smaller, and ∆ increases. The exponent therefore carries information about the behavioral state of the mouse — the amount of information increases with the difference between the resting and moving exponents.

Whereas saline administration preserved the exponents regardless of whether FC changes were accounted for, this was not true for ibogaine (Fig. 3(c)). If changes to FC are unaccounted for, the exponents do change relative to the baseline. For the down-regulated neurons, the exponents are indistinguishable between the resting and moving phases (Fig. 3(c)), suggesting the behavioral state can no longer be inferred from this sub-population. The decreased difference between exponents *τ*_*rst*._ and *τ*_*mv*._ for up-regulated neurons also indicates that this subpopulation no longer carries as much information about behavior. These findings indicate that when changes to FC are accounted for, the information is preserved, but has been routed differently relative to the baseline. Moreover, surrogates that maintain each neurons firing rate and preserve their correlation with the track speed (Sec. S1F) do not exhibit power-law statistics (Fig. S10), implying this is not a simple consequence of the behavioral statistics or sub-population reclassification.

The robustness of the critical brain hypothesis is further supported by the observation that behaviorally conditioned distributions of neuronal avalanche sizes considering *all* neurons preserve the self-similarity under ibogaine as well (Fig. S11). Moreover, neuronal avalanche duration distributions showed the same behaviorally conditioned power-law characteristics as avalanche sizes for behaviorally-related neurons, with varying exponent *α* (Fig. S12, Tables S3, S4). Whereas *τ* and *α* change with behavioral state, the exponent *γ* that characterizes the relationship between neuronal avalanche size and duration (as in the case of behavioral avalanches), does not. We find *γ* ≈ 0.7 (Fig. S13, Tables S3, S4), regardless of the sub-population and whether changes to FC were accounted for. This value and its invariance are consistent with [43]. Most importantly, Fig. 3(d) (see also Fig. S13) confirms that the scaling relation, 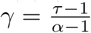 holds, further supporting the critical brain hypothesis [40, 43, 44].

Whereas previous studies have detected changes to, or loss of, signatures of criticality under network changes [7, 10, 28–33], our findings show that psychedelic-induced perturbations to FC preserve the scale-free statistics of neuronal dynamics, one of the signatures of criticality, including those of behaviorally-conditioned sub-populations if rewiring of the FC is accounted for. FC changes under ibogaine also leave behavior and its self-similar features unchanged, as well as some topological features of the functional network. This is despite the loss of information in the RSC encoding of spatial position revealed previously [38]. Thus, our findings suggest that key features of information propagation are invariant and robust against perturbations due to the drug, which alters FC information rerouting.

What is the underlying mechanism for this invariance? It is possible that the drug directly produces the observed changes in FC while preserving criticality, by changing the dynamics (e.g. neural excitability) of individual neurons in the RSC and/or the communication between them, or by changing the inputs to the RSC (consistent with the proposed multi-stability [45]). Alternatively, the invariance could be due to the brains adaptation to the drug-induced perturbation, exhibiting a specific example of self-organized criticality [6], or, some combination of all these mechanisms. For all these mechanisms the structure of the underlying network must allow for sufficiently flexible functional connectivity. Indeed, this flexibility is maximized for efficient information processing in healthy brains operating near criticality [21, 46]. Some have even suggested that the psychedelic state may actually be closer to criticality and allows greater flexibility than the normal waking state [47].

To extend our study of changes in FC and neuronal dynamics to more general contexts, an alternate description of inter-neuron correlations forming a low-dimensional set of neural modes, which span the neuronal activity space [48], is a promising future option. Similarly, any possible connection of our observations to changes in neuronal representations occurring naturally over days and weeks while maintaining output of learned tasks [49, 50] remains to be explored. The mechanism for network reconfiguration over such long time-scales could be qualitatively very different from the drug-induced FC changes on shorter time-scales. However, the degeneracy of network configurations, which is important to maintain behavioral output during representational drift, could also allow for flexibility in FC on shorter time-scales.

In summary, we find ibogaine changes the functional relationship of many neurons with each other and with behavior. However, we found that the scale-free neuronal dynamics are not affected by ibogaine. The critical brain hypothesis indicates this invariance as a necessary condition for the brain to operate near a critical point, wherein the range of stimuli resulting in distinguishable network responses is maximized [5, 18, 51]. Our results imply that FC is variable on time-scales over which the drug affects the system, and suggests that for a given structural arrangement, the functional network adapts to maintain criticality and optimize information propagation in the network. The invariance of neuronal and behavioral dynamics under a potent psychoactive drug that changes FC suggests that these are fundamental dynamical properties robust to perturbations. Finally, our study supports the correspondence between scale-free neuronal and behavioral statistics [36], but whether a direct link between neuronal and behavioral dynamics exists remains an exciting challenge for the future.

## ACKNOWLEDGEMENTS

We would like to thank Adam Neumann for technical support. This work was supported by the Natural Sciences and Engineering Research Council of Canada, New Frontiers Research Fund, Alberta Innovates, Branch Out Neurological Foundation, Izaak Walton Killam Memorial Trusts.

## References

[1] K. D. Harris, Nature Reviews Neuroscience 6, 399 (2005).

[2] J. M. Beggs and D. Plenz, Journal of Neuroscience 23, 11167 (2003).

[3] L. Cocchi, L. L. Gollo, A. Zalesky, and M. Breakspear, Progress in Neurobiology 158, 132 (2017).

[4] J. M. Beggs, The Cortex and the Critical Point: Understanding the Power of Emergence (MIT Press, 2022).

[5] O. Kinouchi and M. Copelli, Nature Physics 2, 348 (2006).

[6] D. R. Chialvo, Nature Physics 6, 744 (2010).

[7] S. H. Gautam, T. T. Hoang, K. McClanahan, S. K. Grady, and W. L. Shew, PLoS Computational Biology 11, e1004576 (2015).

[8] E. D. Fagerholm, G. Scott, W. L. Shew, C. Song, R. Leech, T. Knöpfel, and D. J. Sharp, Cerebral cortex 26, 3945 (2016).

[9] D. Plenz, T. L. Ribeiro, S. R. Miller, P. A. Kells, A. Vakili, and E. L. Capek, Frontiers in Physics 9, 639389 (2021).

[10] T. Bellay, A. Klaus, S. Seshadri, and D. Plenz, Elife 4, e07224 (2015).

[11] D. J. Korchinski, J. G. Orlandi, S.-W. Son, and J. Davidsen, Physical Review X 11, 021059 (2021).

[12] A.-E. Avramiea, A. Masood, H. D. Mansvelder, and K. Linkenkaer-Hansen, Journal of Neuroscience 42, 2221 (2022).

[13] K. Linkenkaer-Hansen, V. V. Nikouline, J. M. Palva, and R. J. Ilmoniemi, Journal of Neuroscience 21, 1370 (2001).

[14] M. Jannesari, A. Saeedi, M. Zare, S. Ortiz-Mantilla, D. Plenz, and A. A. Benasich, Brain Structure and Function 225, 1169 (2020).

[15] J. M. Palva, A. Zhigalov, J. Hirvonen, O. Korhonen, K. Linkenkaer-Hansen, and S. Palva, Proceedings of the National Academy of Sciences 110, 3585 (2013).

[16] Z. Ma, G. G. Turrigiano, R. Wessel, and K. B. Hengen, Neuron 104, 655 (2019).

[17] C. Tetzlaff, S. Okujeni, U. Egert, F. Wörgotter, and M. Butz, PLoS Computational Biology 6, e1001013 (2010).

[18] W. L. Shew, H. Yang, T. Petermann, R. Roy, and D. Plenz, Journal of Neuroscience 29, 15595 (2009).

[19] E. D. Gireesh and D. Plenz, Proceedings of the National Academy of Sciences 105, 7576 (2008).

[20] A. Ponce-Alvarez, A. Jouary, M. Privat, G. Deco, and G. Sumbre, Neuron 100, 1446 (2018).

[21] A. Haimovici, E. Tagliazucchi, P. Balenzuela, and D. R. Chialvo, Physical Review Letters 110, 178101 (2013).

[22] T. Petermann, T. C. Thiagarajan, M. A. Lebedev, M. A. Nicolelis, D. R. Chialvo, and D. Plenz, Proceedings of the National Academy of Sciences 106, 15921 (2009).

[23] G. Hahn, A. Ponce-Alvarez, C. Monier, G. Benvenuti, A. Kumar, F. Chavane, G. Deco, and Y. Frégnac, PLoS Computational Biology 13, e1005543 (2017).

[24] D. Dahmen, S. Grün, M. Diesmann, and M. Helias, Proceedings of the National Academy of Sciences 116, 13051 (2019).

[25] J. Wilting and V. Priesemann, Cerebral Cortex 29, 2759 (2019).

[26] M. G. Kitzbichler, M. L. Smith, S. R. Christensen, and E. Bullmore, PLoS Computational Biology 5, e1000314 (2009).

[27] E. Tagliazucchi, P. Balenzuela, D. Fraiman, and D. R. Chialvo, Frontiers in Physiology 3, 15 (2012).

[28] G. Scott, E. D. Fagerholm, H. Mutoh, R. Leech, D. J. Sharp, W. L. Shew, and T. Knöpfel, Journal of Neuroscience 34, 16611 (2014).

[29] D. Curic, D. Ashby, A. McGirr, and J. Davidsen, Preprint (2023).

[30] M. Yaghoubi, T. de Graaf, J. G. Orlandi, F. Girotto, M. A. Colicos, and J. Davidsen, Scientific Reports 8, 1 (2018).

[31] C. Meisel, A. Storch, S. Hallmeyer-Elgner, E. Bullmore, and T. Gross, PLoS Computational Biology 8, e1002312 (2012).

[32] G. Alamian, T. Lajnef, A. Pascarella, J.-M. Lina, L. Knight, J. Walters, K. D. Singh, and K. Jerbi, Frontiers in Neural Circuits 16 (2022).

[33] P. Sorrentino, R. Rucco, F. Baselice, R. De Micco, A. Tessitore, A. Hillebrand, L. Mandolesi, M. Breakspear, L. L. Gollo, and G. Sorrentino, Scientific Reports 11, 1 (2021).

[34] D. Curic, V. E. Ivan, D. T. Cuesta, I. M. Esteves, M. H. Mohajerani, A. J. Gruber, and J. Davidsen, Journal of Physics: Complexity 2, 045010 (2021).

[35] C. Stringer, M. Pachitariu, N. Steinmetz, C. B. Reddy, M. Carandini, and K. D. Harris, Science 364, eaav7893 (2019).

[36] S. A. Jones, J. H. Barfield, V. K. Norman, and W. L. Shew, Elife 12, e79950 (2023).

[37] P. Garcia-Junco-Clemente, T. Ikrar, E. Tring, X. Xu, D. L. Ringach, and J. T. Trachtenberg, Nature neuroscience 20, 389 (2017).

[38] V. E. Ivan, D. P. Tomàs-Cuesta, I. M. Esteves, D. Curic, M. Mohajerani, B. L. McNaughton, J. Davidsen, and A. J. Gruber, Biological Psychiatry Global Open Science (2023).

[39] D. Plenz and T. C. Thiagarajan, Trends in Neurosciences 30, 101 (2007).

[40] J. Touboul and A. Destexhe, Physical Review E 95, 012413 (2017).

[41] A. Clauset, C. R. Shalizi, and M. E. Newman, SIAM review 51, 661 (2009).

[42] A. Deluca and (2013). Á. Corral, Acta Geophysica 61, 1351

[43] A. J. Fontenele, N. A. de Vasconcelos, T. Feliciano, L. A. Aguiar, C. Soares-Cunha, B. Coimbra, L. Dalla Porta, S. Ribeiro, A. J. Rodrigues, N. Sousa, et al., Physical review letters 122, 208101 (2019).

[44] J. P. Sethna, K. A. Dahmen, and C. R. Myers, Nature 410, 242 (2001).

[45] C. Kirst, M. Timme, and D. Battaglia, Nature communications 7, 11061 (2016).

[46] B. Song, N. Ma, G. Liu, H. Zhang, L. Yu, L. Liu, and J. Zhang, Journal of Neural Engineering 16, 056002 (2019).

[47] R. L. Carhart-Harris, R. Leech, P. J. Hellyer, M. Shanahan, A. Feilding, E. Tagliazucchi, D. R. Chialvo, and D. Nutt, Frontiers in Human Neuroscience, 20 (2014).

[48] J. A. Gallego, M. G. Perich, L. E. Miller, and S. A. Solla, Neuron 94, 978 (2017).

[49] L. N. Driscoll, N. L. Pettit, M. Minderer, S. N. Chettih, and C. D. Harvey, Cell 170, 986 (2017).

[50] M. E. Rule, T. O’Leary, and C. D. Harvey, Current opinion in neurobiology 58, 141 (2019).

[51] D. B. Larremore, W. L. Shew, and J. G. Restrepo, Physical Review Letters 106, 058101 (2011).

