## Supplementary Materials for "Changes in functional connectivity preserve scale-free neuronal and behavioral dynamics"

Anja Rabus[1*], Davor Curic[1], Victorita E. Ivan[2], Ingrid M. Esteves[2], Aaron J. Gruber[2], Jörn Davidsen[1,3†]
*1: Complexity Science Group, Department of Physics and Astronomy*
*University of Calgary, Calgary, Alberta T2N 1N4, Canada*
*2: Canadian Centre for Behavioral Neuroscience*
*University of Lethbridge, Lethbridge, Alberta T1K 3M4, Canada*
*3: Hotchkiss Brain Institute, University of Calgary, Calgary, Alberta T2N 4N1, Canada*
(Dated: September 13, 2023)

### S1. MATERIALS AND METHODS

#### A. Data

Experiments were performed on adult (9-11 month old) Thy1-GCaMP6m mice (n=7) , kept in standard rodent cages at 24°C, during light hours of a 12 h light/dark cycle.

The mice underwent a 5 mm bilateral craniotomy (AP: +1 to -4; ML: -2.5 to +2.5) in the retro-splenial cortex (RSC) and imaging took place through three layers of glass coverslips which covered the lesions using Vetbond. The microscope was held in place by a titanium head plate which was fixed to the skull using dental acrylic.

Mice were water-restricted during training and testing (with daily 30 minutes of free access to water). Weight loss was monitored carefully so as not to exceed 15% of their baseline body weight. Head-fixed mice were trained to move on a non-motorized treadmill belt in daily sessions until they completed at least 20 laps on the belt in 30 minutes. The belt is 4 cm wide and 150 cm long, with four tactile cues placed at different locations. Mice were offered a drop of 10% sucrose solution by a solenoid valve, as a reward after each completed lap.

Continuous neural activity of on average $418 \pm 162$ neurons in the RSC was measured via two-photon (2P) imaging of calcium indicators with 19 Hz scanning rate in 24-minute long recordings. During 2P imaging, the previously trained, head-fixed mice were free to move or stand on the treadmill belt, with the reward still being dispensed after every lap. The microscope images a 835 $\mu m^2$ field of view, with a depth between 135-170 $\mu m$ (layers II/III). An optical encoder measured the rotational motion of the belt, from which the speed of the mice at each time point was obtained. Consecutive baseline-saline recordings exist for all 7 mice, and consecutive baseline-ibogaine recordings exist for 5 of these mice.

*
†

#### B. Classifying populations

Within the RSC, we identify neurons which activate when the animals are moving (resting) as upregulated (downregulated). Correlation values computed between the de-convolved calcium signal of neuronal activity and the track speed form a unimodal distribution. We chose to classify neurons with activity that correlates positively (negatively) with the track speed with $p < 0.05$ as upregulated (downregulated), see Fig. 1(a) for an example and Table S1 for the numbers of classified neurons in the different recordings.

#### C. Functional connectivity

We calculated FC between the de-convolved calcium signal of neurons using $corrcoef$ (MATLAB) and visually represented the connectivity within and between classified populations in adjacency matrices. We were able to characterize changes in FC from baseline to the ibogaine (Fig. 1(b)) and saline (Fig. S1) states, as the locations of individual neurons remain fixed in successive recordings. We quantified the proportion of neurons that switched to, from or between classifications (Fig. 1(c),(d)), calculated the mean absolute distance between matrices (Fig. S2), and determined topological properties of the adjacency matrices (Fig. S3).

#### D. Defining behavioral avalanches

Behavioral avalanches are defined as periods during which the mouse is moving, between periods where the mouse is resting. The avalanche size $l$ is the distance covered by the mouse during the avalanche and the avalanche duration $t$ is the total time that the mouse was in motion during the avalanche. Behavioral avalanches are determined from the track recording, with negative speeds set to zero and a threshold of 0.05 cm/s applied to all recordings, above which the mouse is considered to be moving and below which the mouse is considered to be resting. The same threshold is applied to all mice as the mice are all roughly the same size and weight. All our findings (Figs. 2, S5) are robust with respect to variations in the threshold up to a value of at least 1 cm/s. Results using the latter threshold are shown in Fig. S4, for example. Note that the different distributions of avalanche size, $P(l)$, (Figs. 2(a), S5(a)) and avalanche duration, $P(t)$, (Fig. S5(c)) are obtained from concatenating across all recordings of the same condition, which is reasonable since the effect of ibogaine on the behavioral avalanche distributions is consistent across recordings. It is also necessary in order to ensure a sufficiently high number of behavioral avalanches for reliable estimation of the distributions. Distance $l$ should in theory be limited by reward dispensation every 150 cm. Although there are some instances where the mouse skips the reward and completes more than one lap in one avalanche, in most cases the mice take a break before the end of one lap.

#### E. Defining neuronal avalanches

Neuronal avalanches are traditionally defined as periods of non-zero neuronal activity delineated by periods of inactivity. As total neuronal activity is high relative to 2P scanning rate (19 Hz), periods of quiescence are rare or non-existent. A common approach is to set a threshold — any value in the 2P calcium time series of individual neurons that is below the threshold is set to zero, which generates quiescent periods. As in [1, 2] we use a threshold that maximizes the number of avalanches (or equivalently the avalanche rate) in each recording.

Behaviorally conditional neuronal avalanches are generated as in [2] — periods of rest and movement are identified as described in section S1 D. Specifically, behaviorally conditioned avalanches are generated from up to five avalanches immediately prior to and following a behavioral transition. Avalanches that overlap with the transition, or extend over the last or next moving-to-resting transition are discarded. While one could take all avalanches occurring during the identified rest or movement phases, this was avoided for two reasons: i) While the transition from resting to moving is typically sharp and clear, the reverse transition is not, making it difficult to determine "when" the transition occurred. ii) Once the mouse completes the lap it is given a reward, which often coincided with a movement-to-rest transition and, thus, may result in compounding factors in the neural activity associated with the reward. The total number of behaviorally conditioned neuronal avalanches is closely related to the number of transitions, which did not change substantially under ibogaine, nor did the avalanche rate, as shown in Table S2.

The size of the avalanche, $S$, is the integrated deconvolved calcium signal between periods of quiescence. In order to account for the different number of neurons in each recording, as well as the various levels of activity, we normalize the size by $\Lambda = N\lambda_{\text{avg}}$ [1]. Here, $N$ is the number of neurons within the recording, and $\lambda_{\text{avg}}$ is a measure of the mean activity of the thresholded neuron signal, given by $\lambda_{\text{avg}} = N^{-1}\sum_{i=1}^{N}\lambda_i$, where $\lambda_i$ is the mean number of binary firings of neuron $i$, averaged over the duration of the recording. Figure S9(a) shows the effect of not including this normalization. In all other figures, neuronal avalanches were concatenated across recordings and subjects to maximize the sample size of avalanches under ibogaine, which is limited by the shorter ibogaine recordings (about 10 mins compared to 20 mins for baseline recordings). As follows from Table S2, the overall smaller number of neuronal avalanches for ibogaine prevents us from obtaining reliable estimates for in-subject distributions for ibogaine. We finally note that probability density functions are estimated using logarithmic binning of the concatenated statistics across all relevant recordings of all mice.

The avalanche duration, $T$, is the number of frames that the the avalanche lasted for, converted to seconds by dividing by the sampling rate. Note that because of their strictly discrete nature, logarithmic binning causes visual artifacts in the probability density functions of avalanche durations, in particular for small durations. The distributions in Fig. S12 are instead obtained by frequency counting the number of occurrences of a given duration.

Behaviorally conditioned distributions of neuronal avalanches were calculated separately for up-regulated and down-regulated neurons (Figs. S8, S12). We considered both the case in which changes in FC from pre- to post-administration recordings are accounted for, by reclassifying neurons in the post-administration recording, and the case in which these changes are not accounted for, by using the classification from the pre-administration recording. We also calculated behaviorally conditioned distributions of neuronal avalanche sizes considering *all* neurons, disregarding any classification with respect to correlations with track speed (Fig. S11).

#### F. Surrogates

To directly address the question whether behavioral and neuronal avalanche statistics are trivially related, we conducted a surrogate analysis. Specifically, we randomly permuted neuronal activations for each neuron within the windows prior to and succeeding a moving-to-resting transition. The window is not fixed but rather is set to the time it took for the (up to) five avalanches in the non-surrogate data to occur. This approach to generate surrogates randomises the neuronal firings but (i) preserves each neuron's firing rate during the resting and moving period of each window separately, and (ii) largely preserves the correlation of each neurons with the track speed. This step is repeated five times per transition. Fig. S10 shows the result of this analysis for the up-regulated group in the moving state (where the best fit power-laws are observed) and shows that while the surrogate is capable of generating a wide range of avalanche sizes, the distribution is not a power-law, and indeed no sub-region of the whole distribution could be fit by a power-law, suggesting that the neuronal avalanche statistics we observe are not a direct result of behavioral statistics.

#### G. Power-law Analysis

The analysis of the avalanche distributions was done following [3] and [4]. For both behavioral and neuronal distributions, maximum likelihood estimation (MLE) was performed assuming a power-law defined between a maximum ($s_{max}$) and minimum ($s_{min}$) avalanche size of the form

$$p(\tau|s) = \frac{1-\tau}{s_{max}^{1-\tau} - s_{min}^{1-\tau}} s^{-\tau},$$

for which the log-likelihood function takes the form

$$\ell = \sum_i \log\left(\frac{1-\tau}{s_{max}^{1-\tau} - s_{min}^{1-\tau}} s_i^{-\tau}\right).$$

where $s_i$ are the individual observations. From this, the maximum of $\ell$ as a function of $\tau$ was obtained numerically using the intrinsic MATLAB function *mle*. In order to select $s_{min}$ and $s_{max}$, we first pick an initial (visually guided) pair, perform MLE and perform a Kolmogorov-Smirnov test between $[s_{min}, s_{max}]$ against a theoretical power-law distribution, and calculate the corresponding $p$-value. This is repeated for a new set of $[s_{min}, s_{max}]$. The final domain $[s_{min}, s_{max}]$ is the largest domain that could be obtained that satisfied $p \geq 0.1$. By construction the $p-$value cannot serve as a measure of goodness-of-fit, but is instead the dynamic range of the domain $\Delta = \log_{10}(s_{max}/s_{min})$.

Neuronal avalanche duration distributions exponents were obtained similarly to avalanche sizes, but require a different estimator, as avalanche durations are strictly discrete valued (i.e., in terms of frames) [3],

$$p(\alpha|T) = \frac{T^{-\alpha}}{\zeta(\alpha, T_{min}) - \zeta(\alpha, 1 + T_{max})},$$

where $\zeta(\alpha, T_{min})$ is the Hurwitz zeta function. Examples of the estimated exponents can be found in Fig. S12(a)-(c). Avalanche durations always have a smaller range than avalanche sizes, and thus the fitting range is smaller. However, the exponents typically passed the above established criteria (e.g., p > 0.1) over a fairly large portion of the *available* range.

The exponent $\gamma$, describing the relationship between avalanche size and duration, does not correspond to a probability distribution, thus MLE is not an option. For both neuronal and behavioral avalanches, we first logarithmically binned the avalanche sizes, and calculated the mean duration of avalanches within each bin to obtain $\langle T\rangle(S)$. Robust linear fitting within $[s_{min}, s_{max}]$ (*robustfit* in MATLAB), to the logarithm of the mean avalanche duration for the logarithm of a given size, was used to estimate the slope, which corresponds to $\gamma$. Fig. S13(a) shows an example of this for up-regulated cells in the moving phase, while Fig. S13(b) shows $\gamma$ in the resting and moving phase across all cases tested.

The estimated exponents are summarized in Tables S3 and S4 for the neuronal distributions and in Fig. S5 for the behavioral distributions.

#### H. Scaling Relation

At criticality the exponents $\alpha$, $\tau$, and $\gamma$ are related by the following scaling relation [5]:

$$\gamma = \frac{\tau - 1}{\alpha - 1} \equiv \gamma(\alpha, \tau). \tag{S1}$$

The independent calculation of $\gamma$ and $\gamma(\alpha, \tau)$, and testing how well they agree, is a standard test for criticality [6]. Figure 3(d) shows $\gamma(\alpha, \tau)$ plotted against $\gamma$ from fitting to $\langle T\rangle(S)$ for the up-regulated cells, whereas Fig. S13c shows the same for the down-regulated cells. The uncertainties on $\gamma(\alpha, \tau)$ are obtained via the standard error propagation formula,

$$\delta\gamma^2 = \frac{(\alpha-1)^2}{(\tau-1)^4}\delta\tau^2 + \frac{1}{(\tau-1)^2}\delta\alpha^2.$$


[1] T. Bellay, A. Klaus, S. Seshadri, and D. Plenz, Elife **4**, e07224 (2015).

[2] D. Curic, V. E. Ivan, D. T. Cuesta, I. M. Esteves, M. H. Mohajerani, A. J. Gruber, and J. Davidsen, Journal of Physics: Complexity **2**, 045010 (2021).

[3] A. Clauset, C. R. Shalizi, and M. E. Newman, SIAM review **51**, 661 (2009).

[4] A. Deluca and Á. Corral, Acta Geophysica **61**, 1351 (2013).

[5] J. P. Sethna, K. A. Dahmen, and C. R. Myers, Nature **410**, 242 (2001).

[6] A. J. Fontenele, N. A. de Vasconcelos, T. Feliciano, L. A. Aguiar, C. Soares-Cunha, B. Coimbra, L. Dalla Porta, S. Ribeiro, A. J. Rodrigues, N. Sousa, *et al.*, Physical review letters **122**, 208101 (2019).

| | | Pre | | | Post | | |
|---|---|---|---|---|---|---|---|
| | | Up | Down | Unclassified | Up | Down | Unclassified |
| SALINE | 1 | 222 | 174 | 128 | 200 | 173 | 151 |
| | | 208 | 149 | 103 | 209 | 130 | 121 |
| | 2 | 279 | 196 | 193 | 339 | 113 | 278 |
| | | 344 | 97 | 245 | 305 | 85 | 526 |
| | 3 | 111 | 42 | 42 | 118 | 42 | 35 |
| | | 89 | 33 | 31 | 90 | 29 | 34 |
| | 4 | 118 | 68 | 70 | 110 | 68 | 78 |
| | | 90 | 57 | 65 | 99 | 59 | 54 |
| | 5 | 115 | 40 | 64 | 115 | 58 | 46 |
| | | 95 | 29 | 114 | 103 | 45 | 90 |
| IBOGAINE | 1 | 180 | 188 | 174 | 203 | 157 | 182 |
| | | 236 | 185 | 178 | 217 | 195 | 187 |
| | | 230 | 163 | 191 | 202 | 177 | 205 |
| | 2 | 303 | 112 | 284 | 255 | 168 | 276 |
| | | 269 | 112 | 321 | 183 | 148 | 371 |
| | 3 | 92 | 47 | 222 | 94 | 58 | 209 |
| | | 112 | 78 | 96 | 122 | 75 | 85 |
| | | 154 | 48 | 92 | 136 | 72 | 86 |
| | 4 | 87 | 86 | 113 | 62 | 101 | 123 |
| | | 90 | 75 | 103 | 93 | 83 | 92 |
| | | 102 | 83 | 90 | 102 | 94 | 79 |
| | 5 | 129 | 77 | 238 | 148 | 99 | 197 |
| | | 129 | 82 | 161 | 151 | 59 | 162 |
| | | 111 | 53 | 211 | 131 | 57 | 187 |

TABLE S1. Numbers of up-regulated, down-regulated and unclassified neurons in baseline and post-administration (saline or ibogaine) recordings for the five mice for which ibogaine recordings exist. The total number of neurons (Up + Down + Unclassified) imaged in the pre- and post-administration recordings is the same.

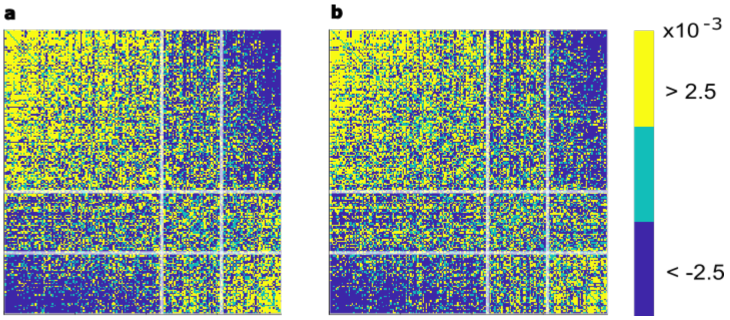


FIG. S1. (a) Correlations between neurons in the baseline recording. Neurons are in order of most correlated to most anti-correlated from left to right and top to bottom, with the white lines separating classified from unclassified groups. (b) Correlations between neurons in the saline recording, preserving the classification of neurons obtained from the baseline recording in (a).

| | Num. Avs | Num. Transitions | Av. Rate ($s^{-1}$) |
|---|---|---|---|
| Baseline | 11895 | 47 ± 17 | 0.22 |
| Saline | 5692 | 47 ± 14 | 0.26 |
| Ibogaine | 2715 | 41 ± 8 | 0.24 |

TABLE S2. The number of avalanches for up-regulated cells for each condition as well as the average number of transitions per recording and avalanche rate. The number of avalanches for down-regulated cells, or when FC changes are not accounted for are roughly the same as this is predominantly determined by the number of transitions.

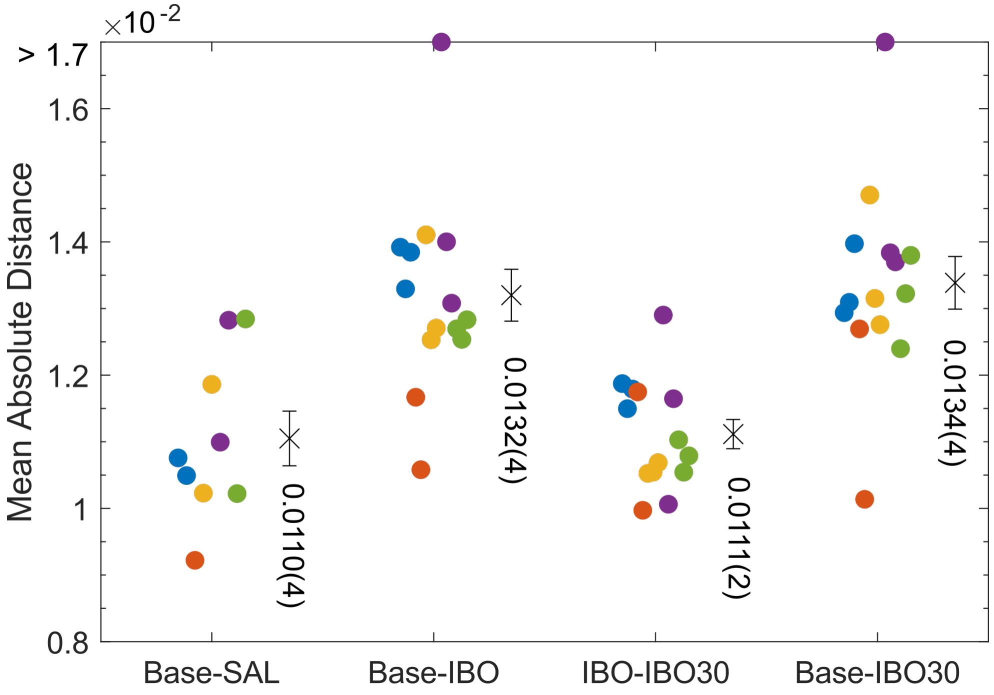


FIG. S2. Mean absolute distance between connectivity matrices. From left to right: distance between saline (SAL) in the baseline (Base) classification and preceding baseline recordings, ibogaine (IBO) in the baseline classification and preceding baseline recordings, 30-minute post-ibogaine (IBO30) in the ibogaine classification and preceding ibogaine recordings, 30-minute post-ibogaine in the baseline classification and preceding baseline recordings. Each color represents an animal. Crosses and errorbars indicate the mean and standard error.

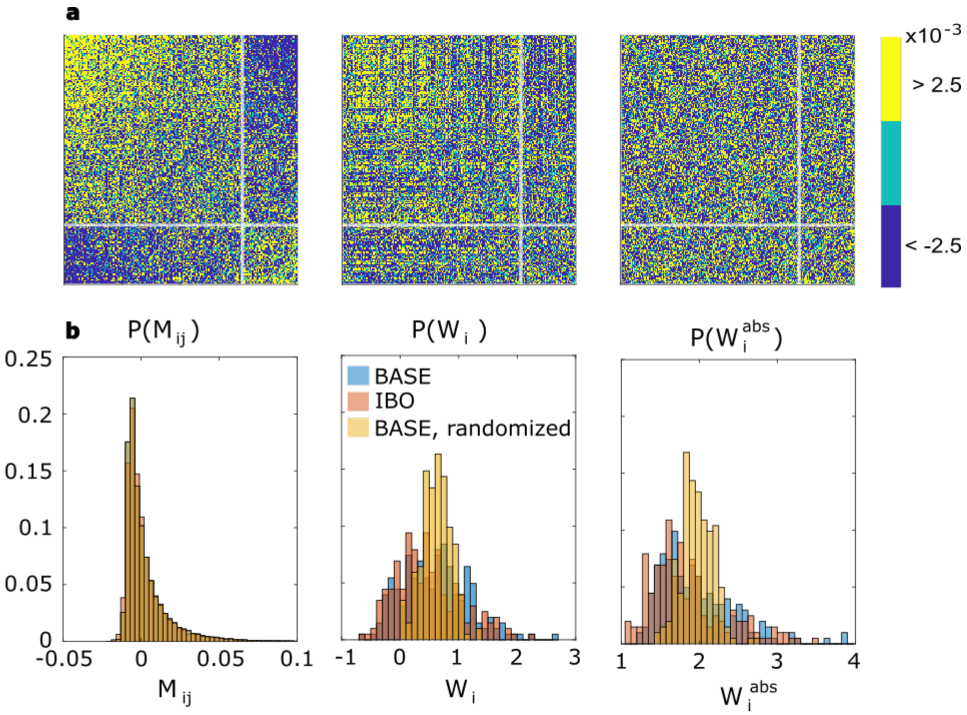


FIG. S3. Representative example recording (a) Left: Connectivity matrix from a representative baseline recording. Middle: Connectivity matrix from the following ibogaine recording. Right: Baseline matrix from the right panel, with entries randomized. (b) Left: Histogram of matrix entries. Middle: Weighted degree distribution. Right: Absolute-valued weighted degree distribution, of the matrices in (a).

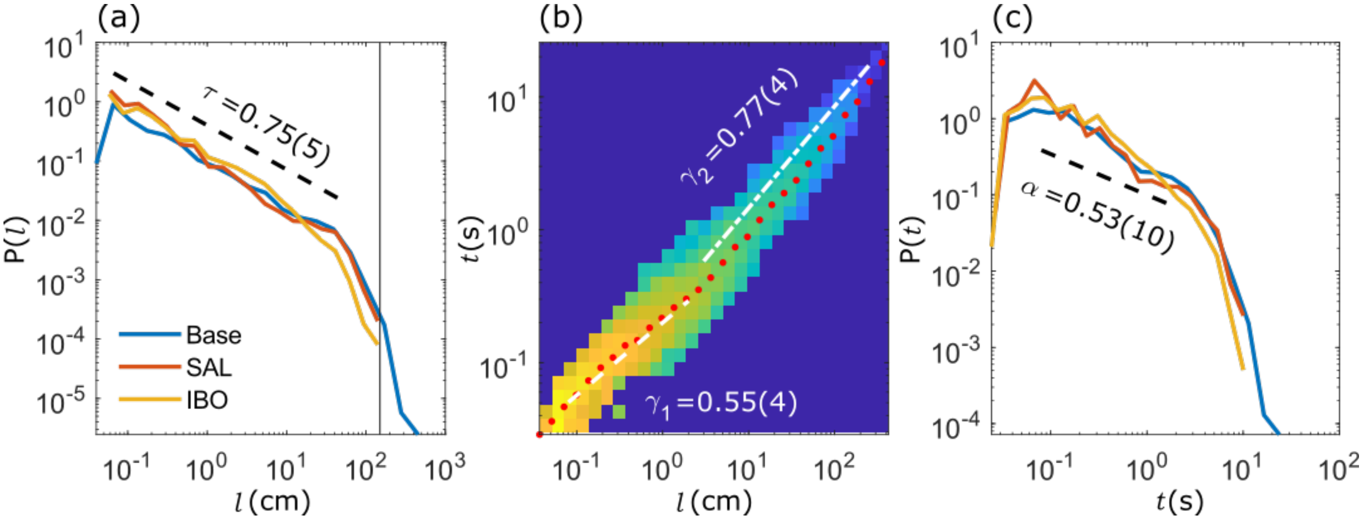


FIG. S4. Scale-free behavioral dynamics when the speed is thresholded at 1 cm/s (see Sec. S1D). (a) Distribution of behavioral avalanche sizes. The exponent $\tau$ was estimated for the baseline recording. The vertical line at 150cm indicates the belt length. (b) Relationship between avalanche duration and size, for all baseline recordings only. Pixel color represents the density of points, and red dots indicate the average $\langle t \rangle$ for a given $l$. The exponents $\gamma_1$ and $\gamma_2$ are given for two fit ranges (white dashed lines). (c) Distribution of behavioral avalanche durations. The exponent $\alpha$ was calculated from $\tau$ in (a) and $\gamma_1$ in (b) using the scaling relation given in Eq. (S1), and plotted for the range of $t$ used to find $\gamma_1$ in (b).

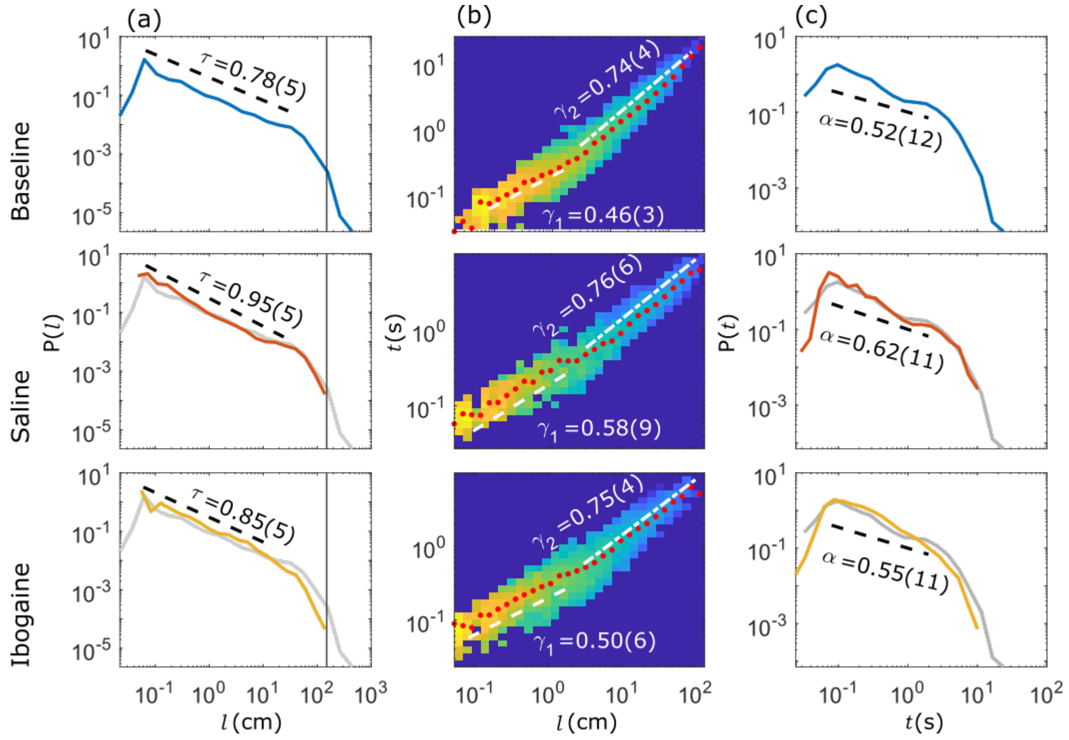


FIG. S5. (a) Behavioral avalanche size distributions for baseline (top), saline (middle) and ibogaine (bottom). Exponents $\tau$ were estimated over two orders of magnitude. (b) Relationship between size and duration for each condition. Exponents $\gamma_1$ and $\gamma_2$ were obtained for separate ranges corresponding to two regimes. (c) Duration distributions for each condition. Exponents $\alpha$ were calculated from $\tau$ in (a) and $\gamma_1$ in (b) using the scaling relation given in Eq. (S1) which must be satisfied at criticality, and plotted over the range of $\gamma_1$.

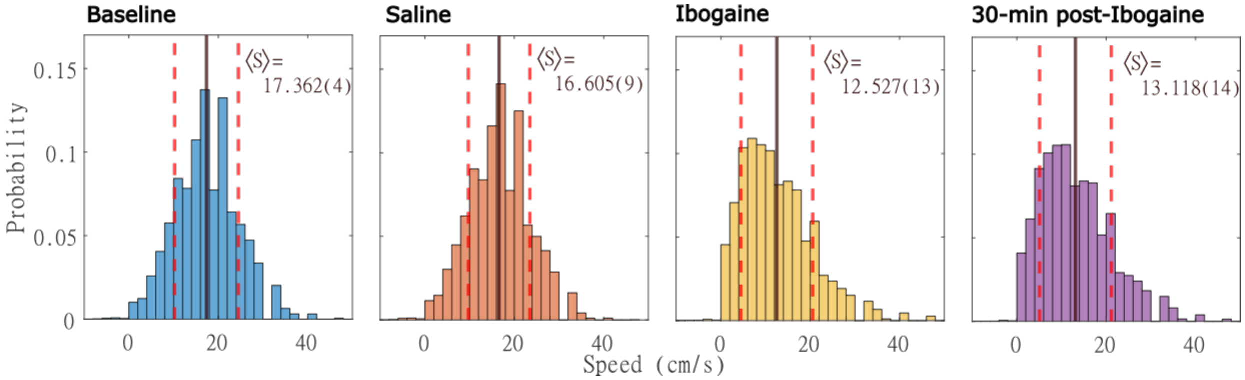


FIG. S6. Normalized distributions of speed for all baseline, saline, ibogaine, and 30-minutes post-ibogaine recordings. Periods where the animals are resting have been excluded. Mean speed $\langle S\rangle$ is given for each distribution (black line), and one standard deviation is indicated (red dashed line).

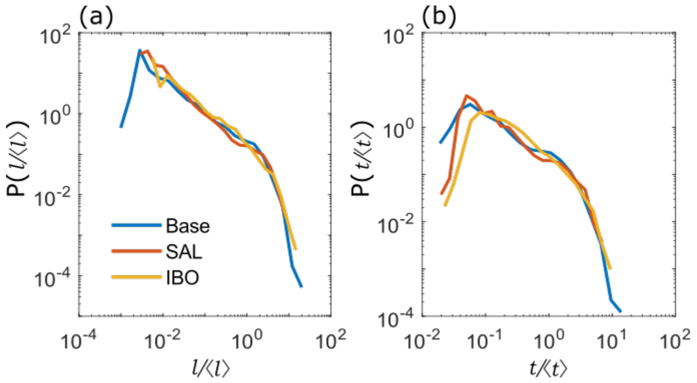


FIG. S7. Dimensionless (a) size and (b) duration distributions for behavioral avalanches for baseline (BASE), saline (SAL) and ibogaine (IBO). The arguments $l$ and $t$ have been re-scaled by averages $\langle l\rangle$ and $\langle t\rangle$ obtained from their respective distributions, which removes the difference in cut-off under ibogaine.

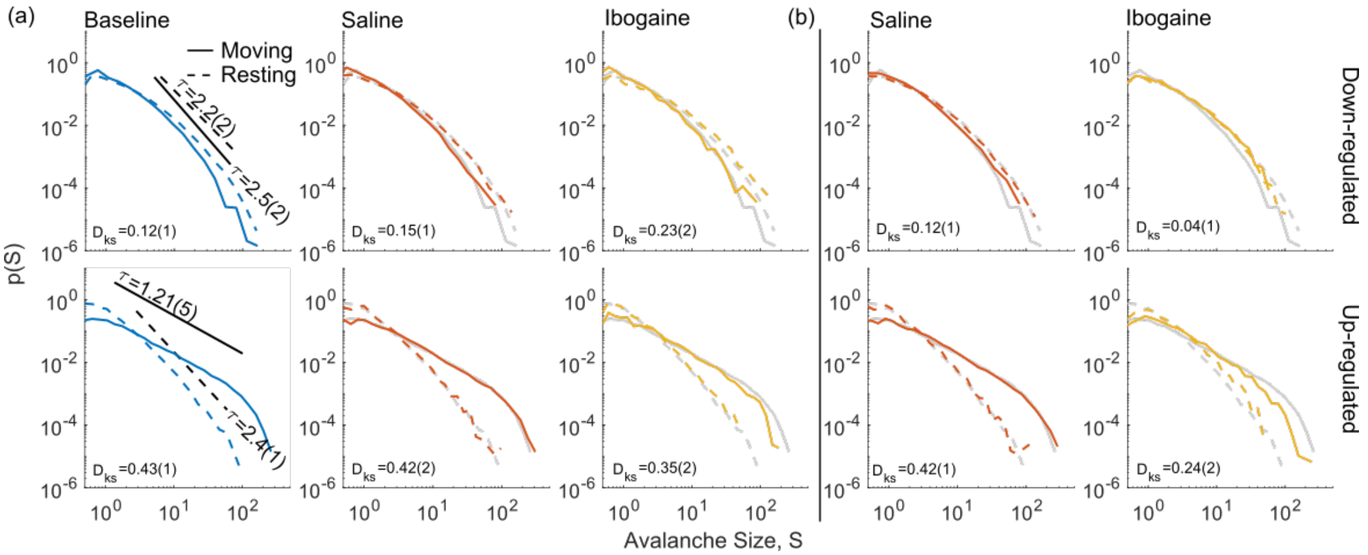


FIG. S8. (a) Behaviorally conditioned neuronal avalanche size distribution with changes in FC are accounted for. In the first column the estimated exponents, represented by the slope of the line, is also shown. The Kolmogorov-Smirnov distance between the resting and moving distributions is also shown ($D_{ks}$). Top row is down-regulated cells and bottom row is up-regulated cells. Solid lines represent the moving (mv.) case and dashed the resting (rst.) case. The gray curves are the same as in the baseline panels and are for visual reference. (b) Same as (a) but when the changes in FC are not accounted for.

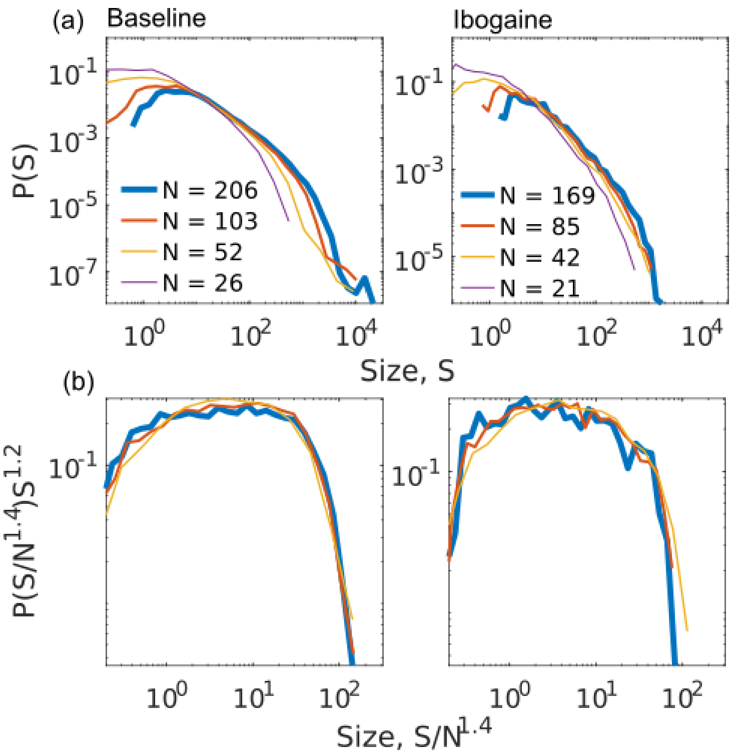


FIG. S9. (a) Movement conditional avalanche size distributions for different numbers of up-regulated neurons without the rescaling factor $\Lambda$ for both baseline (left) and ibogaine (right). (b) Finite scaling analysis shows that subsampling results only in a rescaling of the cut-off, with similar scaling exponents in both baseline and ibogaine.

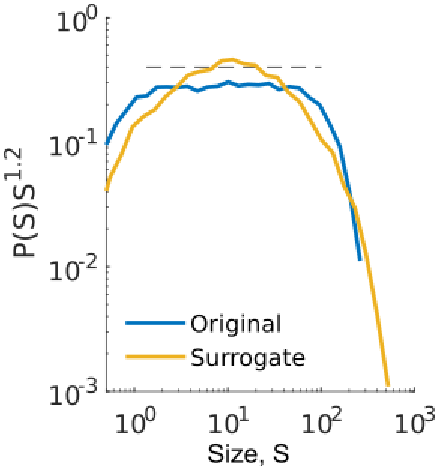


FIG. S10. Surrogate neuronal avalanche size distribution generated by temporal shuffling while maintaining neuron firing rates as described in Section S1 F (yellow), rescaled to the exponent fit to the corresponding non-surrogate avalanches (blue). Dashed line shows fitting domain of the non-surrogate avalanches. No range for power-law fitting was found for the surrogate.

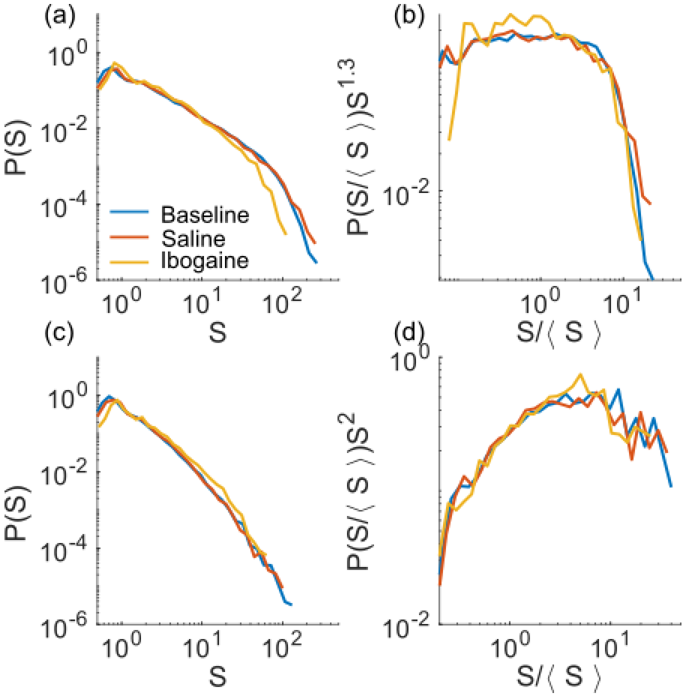


FIG. S11. (a) Avalanche size statistics with no sub-sampling in baseline, saline and ibogaine for the moving phase. (b) Rescaling the distribution by the mean and the critical exponent $\tau = 1.3$ shows curve collapse across all of the three cases. (c) and (d) show the same as (a) and (b) but for the resting state.

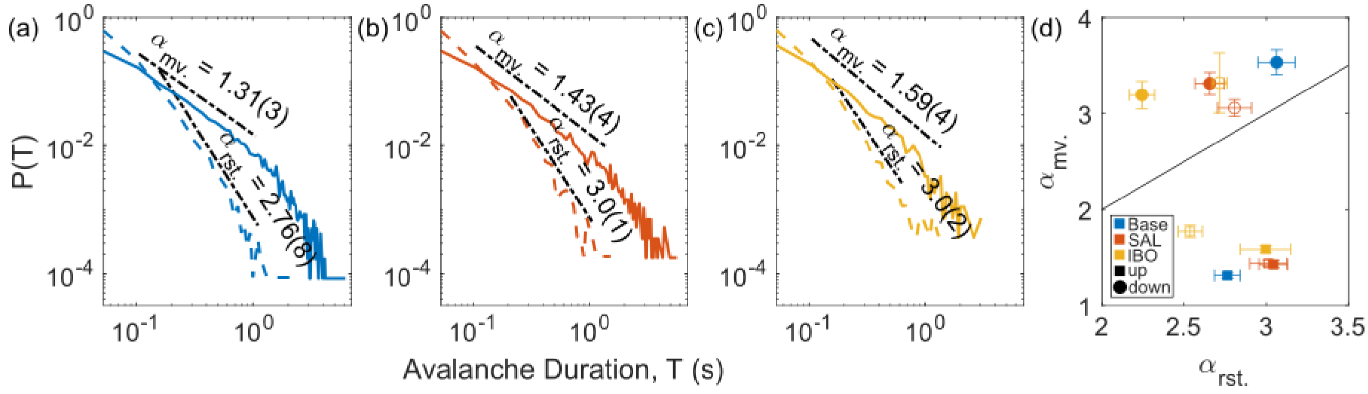


FIG. S12. (a-c) Neuronal avalanche duration distributions for up-regulated cells during movement (solid) and rest (dashed) for baseline, saline, and ibogaine, respectively. Black lines represent region of fitting. (d) The estimated exponents across all cases in the moving case, against the resting case. Solid markers represent the cases wherein changes in FC are accounted for, while hollow markers are the cases in which they are not.

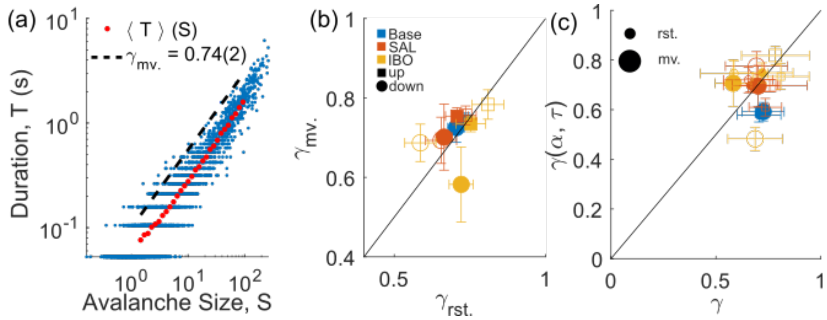


FIG. S13. (a) Baseline example of avalanche duration against size for up-regulated cells in the moving phase. Red dots are the average duration for a given size bin. The slope of the dashed line is $\gamma$. (b) $\gamma$ for moving phase plotted against resting phase. Solid (hollow) symbols represent cases for which changes to functional connectivity have been accounted (unaccounted) for. (c) $\gamma(\alpha, \tau)$ obtained from the scaling relation given in Eq. (S1) plotted vs. the directly estimated value of $\gamma$ from fitting to $\langle T \rangle(S)$ for different conditions for down-regulated cells. Large markers denote moving period, small markers for resting.

| | $\tau_{rst.}$ | $\Delta_{rst.}$ | $\tau_{mv.}$ | $\Delta_{mv.}$ | $\alpha_{rst.}$ | $\Delta_{rst.}$ | $\alpha_{mv.}$ | $\Delta_{mv.}$ | $\gamma(\alpha,\tau)_{rst.}$ | $\gamma_{rst.}$ | $\gamma(\alpha,\tau)_{mv.}$ | $\gamma_{mv.}$ |
|---|---|---|---|---|---|---|---|---|---|---|---|---|
| Baseline down. | 2.2(2) | 1.09 | 2.5(2) | 1.11 | 3.0(1) | 0.77 | 3.5(1) | 0.77 | 0.6(1) | .70(2) | 0.6(1) | .72(4) |
| Saline down. | 2.2(2) | 1.13 | 2.6(2) | 1.13 | 2.7(1) | 0.90 | 3.3(1) | 0.84 | 0.7(1) | .66(3) | 0.7(1) | .70(8) |
| Ibogaine down. | 1.9(2) | 1.23 | 2.5(2) | 1.22 | 2.2(1) | 0.84 | 3.2(1) | 0.84 | 0.74(1) | .72(4) | 0.70(8) | .6(1) |
| Baseline up. | 2.4(1) | 1.32 | 1.21(5) | 1.87 | 2.76(8) | 0.84 | 1.31(2) | 1.0 | 0.77(7) | .73(2) | 0.7(2) | .74(2) |
| Saline up. | 2.3(1) | 1.37 | 1.30(7) | 1.86 | 3.04(8) | 0.70 | 1.43(4) | 1.11 | 0.62(7) | .71(4) | 0.7(2) | .75(2) |
| Ibogaine up. | 2.3(2) | 1.24 | 1.3(1) | 1.78 | 3.0(1) | 0.60 | 1.58(4) | 1.11 | 0.64(9) | .75(5) | 0.7(1) | .73(2) |

TABLE S3. Estimated critical exponents for classified populations for baseline, saline and ibogaine recordings when changes to FC have been accounted for. Exponents are estimated during the resting ($rst.$) and moving phases ($mv.$) of behavior.

| | $\tau_{rst.}$ | $\Delta_{rst.}$ | $\tau_{mv.}$ | $\Delta_{mv.}$ | $\alpha_{rst.}$ | $\Delta_{rst.}$ | $\alpha_{mv.}$ | $\Delta_{mv.}$ | $\gamma(\alpha,\tau)_{rst.}$ | $\gamma_{rst.}$ | $\gamma(\alpha,\tau)_{mv.}$ | $\gamma_{mv.}$ |
|---|---|---|---|---|---|---|---|---|---|---|---|---|
| Saline down. | 2.3(2) | 1.13 | 2.6(3) | 1.10 | 2.8(1) | .77 | 3.06(9) | 1.00 | 0.7(1) | 0.65(5) | 0.7(1) | 0.69(6) |
| Ibogaine down. | 2.3(3) | 1.29 | 2.1(3) | 1.25 | 2.71(5) | .84 | 3.3(3) | 0.77 | 0.74(1) | 0.58(5) | 0.5(1) | .69(5) |
| Saline up. | 2.2(1) | 1.29 | 1.30(7) | 1.67 | 3.0(1) | 1.04 | 1.43(3) | 1.11 | 0.61(7) | 0.73(3) | 0.7(2) | .75(2) |
| Ibogaine up. | 2.1(2) | 1.34 | 1.6(1) | 1.63 | 2.54(7) | 0.90 | 1.77(6) | 1.17 | 0.73(1) | 0.81(5) | .81(2) | .78(4) |

TABLE S4. Estimated critical exponents for classified populations for baseline, saline and ibogaine recordings when changes to FC have *not* been accounted for. Exponents are estimated during the resting ($rst.$) and moving phases ($mv.$) of behavior.
